# A Novel Small Animal Model of Norovirus Diarrhea

**DOI:** 10.1101/2020.03.19.999185

**Authors:** Alexa N. Roth, Emily W. Helm, Carmen Mirabelli, Erin Kirsche, Jonathan C. Smith, Laura B. Eurell, Sourish Ghosh, Nihal Altan-Bonnet, Christiane E. Wobus, Stephanie M. Karst

## Abstract

Human noroviruses are the leading cause of severe childhood diarrhea worldwide yet we know very little about their pathogenic mechanisms. Murine noroviruses cause diarrhea in interferon-deficient adult mice but these hosts also develop systemic pathology and lethality, reducing confidence in the translatability of findings to human norovirus disease. Herein we report that a murine norovirus causes self-resolving diarrhea in the absence of systemic disease in wild-type neonatal mice, thus mirroring the key features of human norovirus disease and representing a robust norovirus small animal disease model. Intriguingly, lymphocytes are critical for controlling acute norovirus replication while simultaneously contributing to disease severity, likely reflecting their dual role as targets of viral infection and key components of the host response.

## MAIN TEXT

Human noroviruses are the leading cause of severe childhood diarrhea and gastroenteritis outbreaks worldwide^1–3^ yet we know very little about their pathogenic mechanisms. While human noroviruses cause modest diarrhea in gnotobiotic piglets, gnotobiotic calves, and miniature piglets^4–6^, a limiting factor in studying norovirus pathogenesis is the lack of tractable small animal models that recapitulate key features of disease observed in infected individuals. The first murine norovirus, MNV-1, was discovered nearly two decades ago^7^ and many other MNV strains have been reported since then. MNV strains segregate into two phenotypic categories which differ in many aspects of pathogenesis, including their rate of clearance from infected hosts and cell tropism. MNV-1 is the prototype acute strain. It infects immune cells in the gut-associated lymphoid tissue (GALT) and reaches peak intestinal titers 1-2 days post-infection (dpi) with clearance from the host by 7-14 dpi^8^. MNV-3 and MNV-CR6 are commonly referred to as persistent strains since they establish a life-long colonic infection^9–11^. The persistent reservoir is a rare type of intestinal epithelial cell, the tuft cell^12^. While major strides have been made in understanding the cellular and tissue tropism of noroviruses using the MNV model system, a limitation of the murine model is the lack of diarrhea in wild-type laboratory mice infected with any of the MNV strains studied to date^13–15^. While MNV-1 does not cause overt disease in adult wild-type mice, infection of interferon (IFN)-deficient *Stat1*^*-/-*^ and *IFNαβγR*^*-/-*^ adult mice causes severe weight loss and diarrhea similar to human norovirus-infected individuals^13–16^. The persistent strains MNV-3 and MNV-CR6 cause less disease than MNV-1 in IFN-deficient mouse strains^14,17^. We confirmed this virulence profile by infecting adult wild-type or *Ifnar1*^*-/-*^ mice with MNV-1, MNV-3, or MNV-CR6. While MNV-1 caused weight loss and fatality of *Ifnar1*^*-/-*^ mice by 4 dpi, MNV-3 and MNV-CR6 failed to cause overt disease **(Fig. 1a-b).** Overall, these results indicate that MNV can cause disease in adult mice but virulence is regulated by an IFN response and viral genetic determinants. Moreover, MNV-1 infection of IFN-deficient mice causes systemic disease and mortality in addition to diarrhea which are not hallmarks of human norovirus infection. Thus, we sought to develop a more relevant disease model.

**Figure 1.**
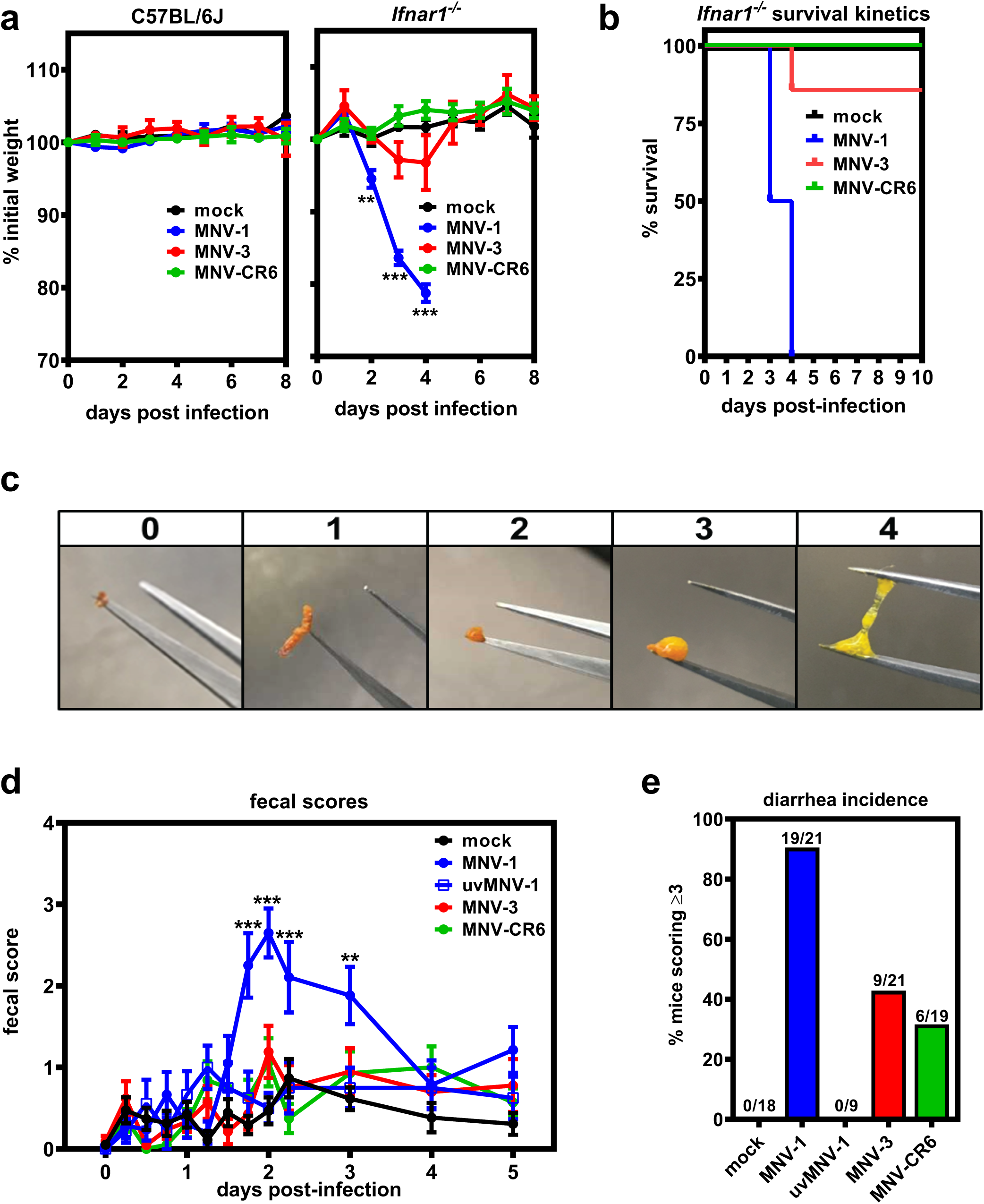
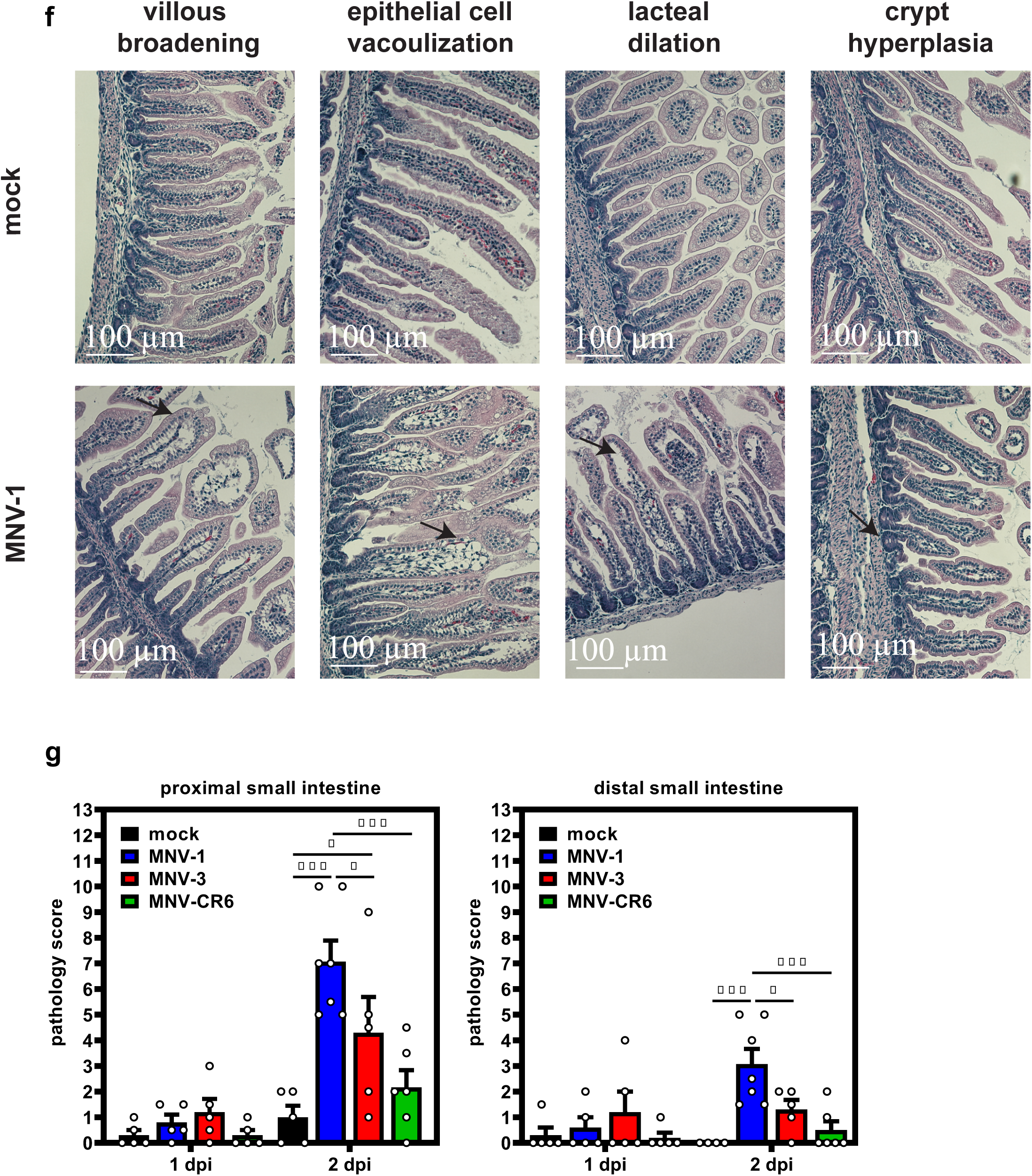
MNV infection causes diarrhea in neonatal mice in a virus strain-dependent manner. **a-b)** Groups of C57BL/6J (*n =* 6 for mock, *n =* 8 for MNV-1, *n =* 6 for MNV-3, *n =* 8 for MNV-CR6) or C57BL/6J-*Ifnar*^*-/-*^ (*n =* 6 for mock, *n =* 8 for MNV-1, *n =* 7 for MNV-3, *n =* 7 for MNV-CR6) mice were perorally infected with 10^7^ TCID_50_ units of MNV-1, MNV-3, MNV-CR6 or mock inoculum and followed for weight loss **(a)** and survival **(b)**. **c)** Representative images of fecal samples indicating each score used to assess for diarrhea are shown. **d-e)** Groups of 3-day old BALB/c pups were infected with 10^8^ TCID_50_ units of MNV-1, UV-inactivated MNV-1, MNV-3, MNV-CR6 or mock inoculum by oral gavage. Pups (*n =* 19 for mock, *n =* 21 for MNV-1, *n =* 9 for UV-inactivated MNV-1, *n =* 21 for MNV-3, *n =* 19 for MNV-CR6) were monitored for fecal consistency by palpating their abdomens **(d)**. The proportion of mice scoring a 3 or 4 at any time point over the 5 d course of infection is presented as the incidence of diarrhea **(e)**. **f-g)** Small intestinal sections collected from neonates infected with 10^8^ TCID_50_ units of MNV-1, MNV-3, MNV-CR6, or mock inoculum at 1 dpi (*n =* 5 for mock, *n =* 5 for MNV-1, *n =* 5 for MNV-3, *n =* 5 for MNV-CR6) and 2 dpi (*n =* 5 for mock, *n =* 7 for MNV-1, *n =* 5 for MNV-3, *n =* 6 for MNV-CR6) were stained with hematoxylin and eosin. Representative images of notable pathology are shown **(f)**. Sections were scored blindly by an animal veterinarian for pathological changes **(g)**. Error bars denote standard errors of mean in all figures. *P* values were determined using two-way ANOVA with corrections for multiple comparisons. (*P < 0.05, **P < 0.01, ***P < 0.001).

Like noroviruses, rotaviruses cause severe acute diarrhea in people yet fail to cause disease in adult wild-type mice. However, it is well-established that neonatal wild-type mice are susceptible to rotavirus-induced diarrhea^18–20^. Moreover, disease severity following human norovirus infection is greater in younger children^21–23^. Thus, we tested whether MNV causes disease in genetically wild-type neonatal mice. Indeed, oral inoculation of 3-d old BALB/c mice with acute MNV-1 caused diarrhea beginning at 42 hours post-infection (hpi) and resolving by 4 dpi **(Fig. 1c-d**, **and Supplementary Fig. 1a)**, with a 91% incidence **(Fig. 1e)**. Virus replication was required for diarrhea induction, as demonstrated by the absence of disease in pups inoculated with UV-inactivated MNV-1 **(Fig. 1d-e)**. Furthermore, MNV-1 diarrhea occurred in a dose-dependent manner **(Supplementary Fig. 1b)**. The incidence of diarrhea was significantly higher in MNV-1-infected 3-d old BALB/c mice compared to 4-d old BALB/c mice, with a 91% versus 49% incidence, respectively **(Supplementary Fig. 2a)**, and slightly, although not significantly, higher in female BALB/c mice than in male mice **(Supplementary Fig. 2b)**. MNV-1 induced diarrhea in BALB/c mice at two independent research institutions (University of Florida and the National Institutes of Health). For all other experiments described herein, we used 3-d old BALB/c mice of mixed sex. We next tested whether the persistent strains MNV-3 and MNV-CR6 likewise caused diarrhea in neonatal mice. Compared to the 91% incidence of diarrhea in MNV-1-infected BALB/c pups, MNV-3 and MNV-CR6 caused diarrhea in 43% and 32% of pups, respectively **(Fig. 1d-e)**. Collectively, these data reveal that MNV-1 infection causes self-resolving acute diarrhea in neonatal BALB/c mice and that MNV-3 and MNV-CR6 cause a reduced incidence of diarrhea relative to MNV-1, mirroring their virulence patterns previously observed in adult IFN-deficient mice^14,17^.

We next determined whether MNV-induced diarrhea is associated with histopathological changes along the intestinal tract. A number of human volunteer studies have been carried out where intestinal biopsies were obtained from subjects at the time of norovirus symptoms^24–29^: While the epithelium remained intact with no gross lesions and there was only a minimal to moderate increase in lamina propria cellularity noted in these patients, there were consistent infection-associated pathologies including villous blunting and broadening, vacuolated and disorganized epithelial cells, and crypt cell hyperplasia. Histopathological changes in MNV-infected neonatal mice are entirely consistent with these human biopsies: While the epithelium itself remained intact and only modest inflammation was observed, there was villous broadening, epithelial disorganization and vacuolization, and crypt cell hyperplasia **(Fig. 1f and Supplementary Fig. 3)**. We also observed lacteal dilation, a pathology previously noted in MNV-4-infected *Stat1*^*-/-*^ mice^30^ that can be indicative of impaired fat absorption. Overall pathology was more pronounced in the proximal half of the small intestine than the distal half, and it positively correlated with diarrhea incidence when comparing intestinal sections from pups infected with MNV-1, MNV-3, and MNV-CR6 **(Fig. 1g and Supplementary Fig. 3)**.

Because MNV-1 caused the highest incidence of diarrhea in neonates, we further characterized its pathogenesis in this model. Virus titers were similar in all regions of the gastrointestinal tract at the peak of disease, 2 dpi, and splenic titers were high at this time point **(Fig. 2a)**. In order to confirm viral replication in vivo, we infected neonatal mice with light-sensitive neutral red-labeled MNV-1 which enables differentiation between input and newly synthesized virus^31,32^. Newly synthesized virus was detectable in all tissues as early as 0.5 dpi and peaked at 1 dpi **(Fig. 2b)**. We next examined the cell tropism of MNV-1 during symptomatic infection. We previously showed that MNV-1 infects immune cells in the GALT of the distal small intestine of wild-type adult mice^8^. To determine whether there is a similar tropism during symptomatic norovirus infection, intestinal sections from MNV-1-infected neonates were hybridized with probes to the viral positive- and negative-sense RNA species and analyzed by RNAscope-based in situ hybridization. At 1 dpi, the peak of viral replication, viral positive-sense RNA was detected in both epithelial and subepithelial cells in the intestinal villi and GALT but negative-sense RNA indicative of viral replication was exclusively detected in subepithelial cells in the GALT and the lamina propria **(Fig. 2c)**. The presence of viral positive-sense, but not negative-sense, viral RNA in intestinal epithelial cells suggests that either virions can enter these cells but not replicate; or that replication occurs in these cells at a level below the limit of detection of our assay. Substantial viral replication was detected in splenocytes, consistent with high splenic titers and with a predominantly immune cell tropism. At 2 dpi, the peak of diarrhea, there was still viral positive-sense RNA along the gastrointestinal tract but minimal negative-sense viral RNA, demonstrating that viral replication in the gut has primarily been controlled by this time point **(Fig. 2d)**. The pattern of viral positive-sense RNA at this time point was variable among the 7 pups analyzed: Four pups contained positive-sense RNA primarily in subepithelial cells of the GALT and lamina propria similar to 1 dpi **(Fig. 2d**, **Pattern 1)** whereas two pups contained positive-sense viral RNA primarily in villous epithelial cells **(Fig. 2d**, **Pattern 2)**. Intriguingly, this signal was confined to the distal region of the small intestine and to the upper half of the villi in this region. The remaining pup displayed an intermediate phenotype, with positive-sense RNA detected in subepithelial and epithelial cells. All pups had substantial positive-sense RNA and variable amounts of negative-sense RNA in the spleen at 2 dpi. Overall, viral replication was observed in subepithelial cells in the intestine and spleen.

**Figure 2.**
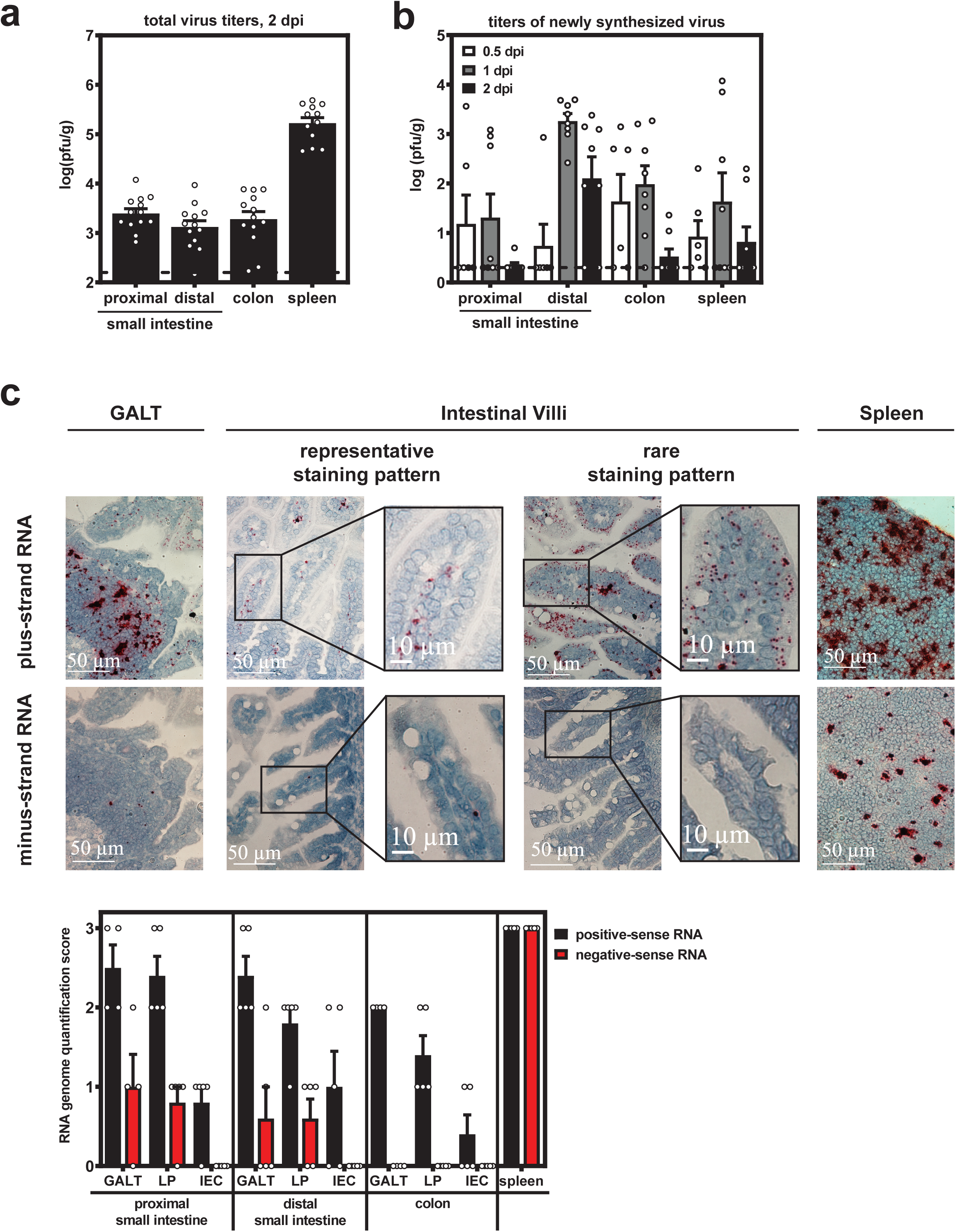
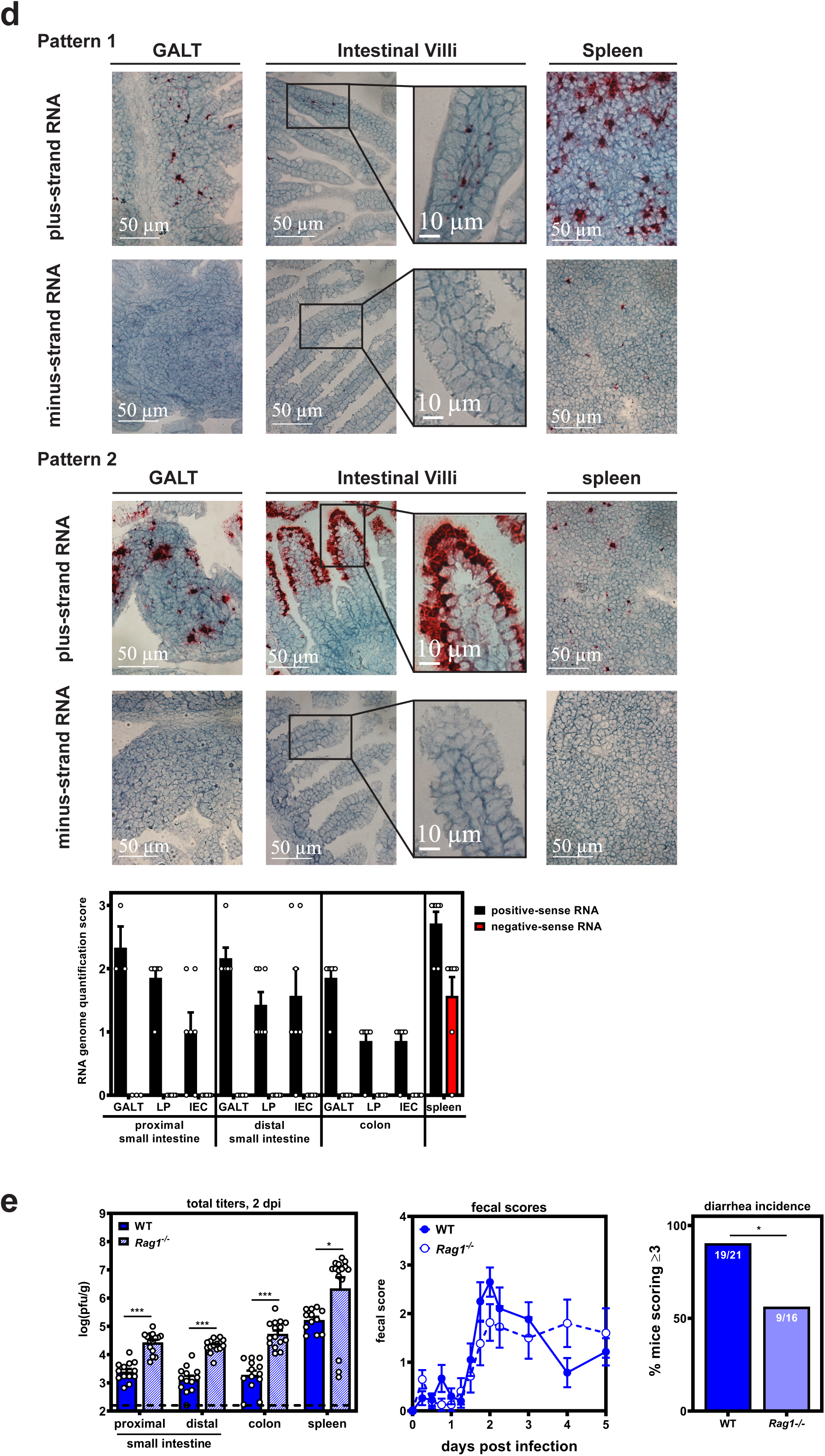
MNV-1 replicates in intestinal subepithelial cells and splenocytes during symptomatic infection. **a)** Three-day old BALB/c pups (*n =* 13) were infected with 10^8^ TCID_50_ units of MNV-1. At 2 dpi, viral titers were determined in the indicated segments of the intestinal tract and spleen by plaque assay. **b)** Three-day old BALB/c pups were infected with 2.5 x 10^4^ pfu of neutral red-labeled MNV-1 by oral gavage At 0.5 dpi (*n* = 5), 1 dpi (*n =* 8) and 2 dpi (*n =* 8), plaque assays were used to measure newly synthesized virus. **c-d)** Tissue sections from mock-inoculated or MNV-1-infected mice collected at 1 dpi **(c;** *n* = 5 per group) and 2 dpi **(d;** *n* = 7 per group) were probed for viral positive-sense and negative-sense RNA. Representative images of gut-associated lymphoid tissue (GALT), intestinal villi, and splenic tissue are shown. The amount of positive-sense and negative-sense viral RNA detected was scored for each pup, as described in the methods. **e)** Groups of 3-day old BALB/c-*Rag1*^*-/-*^ pups were infected with 10^8^ TCID_50_ units MNV-1 by oral gavage. At 2 dpi (*n =* 15), total viral titers were determined in the indicated segments of the intestinal tract and spleen by plaque assay and compared to those of wild-type BALB/c pups. Additional BALB/c-*Rag1*^*-/-*^ pups (*n* = 16) were infected with 10^8^ TCID_50_ units MNV-1 by oral gavage and monitored for fecal consistency over a 5-day time course by palpating their abdomens. Fecal scores and diarrhea incidence were compared to those in wild-type BALB/c pups. Error bars denote standard errors of mean in all figures. *P* values were determined using two-way ANOVA with corrections for multiple comparisons. (*P < 0.05, **P < 0.01, ***P < 0.001).

To begin dissecting the role of specific cellular targets in norovirus disease, we infected lymphocyte-deficient *Rag1*^*-/-*^ neonates with MNV-1. Surprisingly, virus titers at 2 dpi were significantly higher in *Rag1*^*-/-*^ mice compared to wild-type controls **(Fig. 2e)**. In spite of these increased virus titers, the incidence of diarrhea was reduced in *Rag1*^*-/-*^ mice (56%) compared to wild-type mice (90%) **(Fig. 2e)**. These collective results suggest that lymphocytes play opposing roles during acute norovirus infection, being important for control of MNV-1 replication while also contributing to disease severity possibly related to their permissiveness to the virus.

The establishment of a small animal model to study norovirus-induced diarrhea in a genetically wild-type host that recapitulates key features of human disease will revolutionize the field’s ability to understand the precise mechanisms underlying norovirus virulence and to rationally design antiviral therapies. Strengths of this new model compared to adult IFN-deficient mice are that disease is limited to the intestinal tract and self-resolving. Disease severity in this model is (i) regulated by viral genetics, (ii) age-dependent, (iii) not associated with disruption of the intestinal epithelium or notable inflammation, and (iv) associated with immune cell infection. In particular, the finding that highly genetically related MNV strains display differences in diarrhea severity will facilitate identification of viral virulence determinants. Underscoring the potential of this new model system, the discovery that neonatal mice are susceptible to rotavirus diarrhea was key to defining viral disease mechanisms such as the roles of rotavirus NSP4 as a viral enterotoxin^33^ and rotavirus stimulation of the enteric nervous system leading to increased intestinal transit associated with diarrhea^34,35^. This newly described model system holds a similar promise to reveal new insights into norovirus disease mechanisms.

## ACKNOWLEDGEMENTS

S.M.K. was funded by NIH R01AI116892, NIH R01AI081921, and NIH R01AI141478. A.N.R was supported by NIH 5T32AI007110. E.W.H was supported by NIH T90DE021990.

## METHODS

### Cells and viruses

Cell lines used in this study tested negative for mycoplasma contamination and were not authenticated. The RAW 264.7 cell line (ATCC) was maintained in Dulbecco’s modified Eagle medium (DMEM; Fisher Scientific) supplemented with 10% fetal bovine serum (FBS, Atlanta Biologicals), 100 U penicillin/mL, and 100 μg/mL streptomycin. Stocks of recombinant MNV-1.CW3 (GenBank accession number KC782764, referred to as MNV-1), MNV-3 (GenBank accession number KC792553), and MNV-CR6 (GenBank accession number JQ237823.1) were generated as described previously^**36**^. Briefly, 10^6^ 293T cells were transfected with 5 µg infectious clone using Lipofectamine 2000 (Life Technologies), cells were freeze-thawed after 1 d, and lysates titrated with a standard TCID_50_ assay^**11**^ as described below and applied to RAW 264.7 cells. RAW 264.7 lysates were freeze-thawed when cultures displayed 90% cytopathic effect and supernatants were clarified by low-speed centrifugation followed by purification through a 25% sucrose cushion. The viral genomes of stocks were sequenced completely to confirm no mutations arose during stock generation and tittered by a standard TCID_50_ assay^**11**^. For the TCID_50_ assay, 8 replicates each of multiple dilutions per sample were applied to RAW 264.7 cells and cytopathicity scored at 7 dpi. Three independent experiments were averaged to determine stock titers. A mock inoculum stock was prepared using RAW 264.7 lysate from uninfected cultures. A neutral red-labeled MNV-1 stock was generated as previously described^**31,37**^. Briefly, RAW 264.7 cells were infected at MOI 0.05 with MNV-1 in the presence of 10 μg/ml neutral red (Sigma), a light sensitive dye. After 2 d, supernatant was collected in a darkened room using a red photolight (Premier OMNI). Stocks were freeze-thawed twice and stored in a light safe box at -80°C. Viral titer was determined by neutral red virus plaque assay without white light exposure. For UV-inactivation of MNV-1, MNV-1 was exposed to 250,000 µJ cm^-2^ UV for 30 min. UV inactivation was confirmed using TCID_50_ assay.

### Mice

Specific-pathogen-free (SPF) mice used in this study were bred and housed in animal facilities at the University of Florida, National Institutes of Health, and University of Michigan. All animal experiments were performed in strict accordance with federal and university guidelines. The animal protocols were approved by the Institutional Animal Care and Use Committee at the University of Florida (study number 20190632), National Institutes of Health (study number H-0299), and University of Michigan (study number 00006658). Experiments were not performed in a blinded fashion nor was randomization used. Sample sizes were determined using power calculations based on our prior experience with the MNV system. Age- and sex-matched adult C57BL/6J (Jackson no. 000664) and C57BL/6J-*Ifnar1*^*-/-*^ (Jackson no. 010830) mice were used in adult mouse infections. Litters of 3- or 4-day old and BALB/c (Charles River no. 028) and 3-day old BALB/c-*Rag1*^*-/-*^ (Jackson no. 003145) pups including males and females were used in neonatal mouse infections. At least 2 independent litters were infected per condition in each experiment.

### Mouse infections and other in vivo procedures

For all adult MNV infection experiments, six- to twelve-week old, sex-matched mice were inoculated perorally (p.o.) with 10^7^ TCID_50_ units of the indicated virus strain. For virulence assays in adult mice, C57BL/6J and C57BL/6J-*Ifnar1-/-* mice were observed for weight loss and survival daily for up to 14 days. The daily weights were compared to day 0 weights to calculate the relative weight loss. For all neonatal mouse experiments, BALB/c or BALB/c-*Rag1*^*-/-*^ mice were infected at 3 or 4 days of age with the indicated dose of the indicated virus strain in a volume of 40μl delivered by oral gavage using a 22ga plastic feeding tube. Mice were observed every 6 h for the first 54 hours post-infection (hpi), then daily thereafter. At each time point, the abdomen of each mouse was palpated to induce defecation. Fecal condition was assessed based on color and consistency according to a 5-point scale: 0, no defecation or solid; 1, firm, orange, does not smear; 2, pasty, orange or mixed color, does not smear; 3, orange or yellow, semi-liquid and smears; 4, yellow, liquid, and smears. For the purpose of calculating the incidence of diarrhea, any pup that received a score of 3 or 4 at any time point was considered positive for disease. For MNV titer determination, tissue samples were harvested and titrated by plaque assay, as previously described^**14,38**^. In certain experiments, neutral red-labeled virus was used in order to differentiate between inoculum and newly synthesized virus along the intestinal tract^**31,37**^. When using tissues from mice infected with neutral red virus, samples were exposed to white light for 30 min prior to titration in order to inactivate input virus.

### Histology

Small and large intestinal tissue sections as well as spleen sections were collected from 3-day old pups infected with mock inoculum or 10^8^ TCID_50_ units MNV-1, MNV-3, or MNV-CR6 at 1 and 2 dpi. Intestinal issues were swiss-rolled and fixed in 10% buffered formalin for 16 hours, transferred to PBS, and then paraffin-embedded and sectioned by the University of Florida Molecular Pathology Core. Serial sections of 4µm thickness were cut from each tissue and slides stained with hematoxylin and eosin by the University of Florida Molecular Pathology Core. Tissue sections were imaged using a Nikon Ti-E widefield microscope with a Nikon DS-Fi2 color camera and NIS Elements software at the University of Florida Cell and Tissue Analysis Core. Histology slides were scored blindly by an animal veterinarian based on an established scoring system^**39**^.

Three categories - inflammatory cell infiltrate, epithelial changes, and mucosal architecture - were scored for two criteria each: Inflammatory cell infiltrate was scored for severity (0 = none; 1 = minimal; 2 = mild with scattered polymorphonuclear neutrophils; 3 = moderate) and extent (0 = none; 1 = mucosal; 2 = mucosal and submucosal). Epithelial changes were scored for crypt hyperplasia (0 = 2-3mm crypt height at 10X; 1 = 3-4mm crypt height; 2 = 4-5mm crypt height; and 3 = ≥5 mm crypt height) and reduction in goblet cells (0 = no reduction; 1 = minimal reduction). Mucosal architecture was scored for vacuolated epithelium (0 = none; 1 = present at villous tip; 2 = present and disorganized, seen at the base) and villous tip swelling (0 = none; 1 = lacteal separation with cellular material in the center; 2 = lacteal dilation). Data are presented as the total pathology score for all criteria per mouse averaged per group **(Fig. 1g)** and separated by category **(Supplementary Fig. 3)**.

### RNAscope ISH

RNAscope ISH assays were performed using the RNAscope 2.5 HD Assay-RED kit according to manufacturer’s instructions (Advanced Cell Diagnostics, Newark, CA). Formalin-fixed, paraffin embedded serial sections of intestinal swiss rolls from mock- and MNV-1-infected pups were hybridized with custom-designed probes targeting positive-sense or negative-sense MNV-1 RNA prior to probe amplification and detection, as we recently described for adult mouse intestinal sections^8^. Sections were counterstained with 50% hematoxylin to visualize tissue morphology and imaged using a Nikon DS-Fi2 color camera and NIS Elements software at the University of Florida Cell and Tissue Analysis Core. Sections were scored based on amount of virus present in the GALT, intestinal lamina propria, and intestinal epithelial cells on a scale from 0 to 3 (0 = 0-1 dots, 1 = rare staining but at least 2 dots, 2 = consistent, distinguishable dots throughout the section, 3 = intense staining with overlapping dots). Positive (PPIB) and negative (DapB) control probes were stained in parallel for all experiments, and six mock mice were stained.

### Statistical Analyses

All data were analyzed with GraphPad Prism software. Error bars denote standard errors of mean in all figures and *P* values were determined using one- or two-way ANOVA with corrections for multiple comparisons. In the case of survival curves and incidence of diarrhea, statistical significance was determined using a Mantel-Cox test (\**P* < 0.05, \*\**P* < 0.01, \*\*\**P* < 0.001).

## DATA AVAILABILITY

All data that support the findings of this study are available from the corresponding author on request.

## FIGURE LEGENDS

**Supplementary Figure 1.**
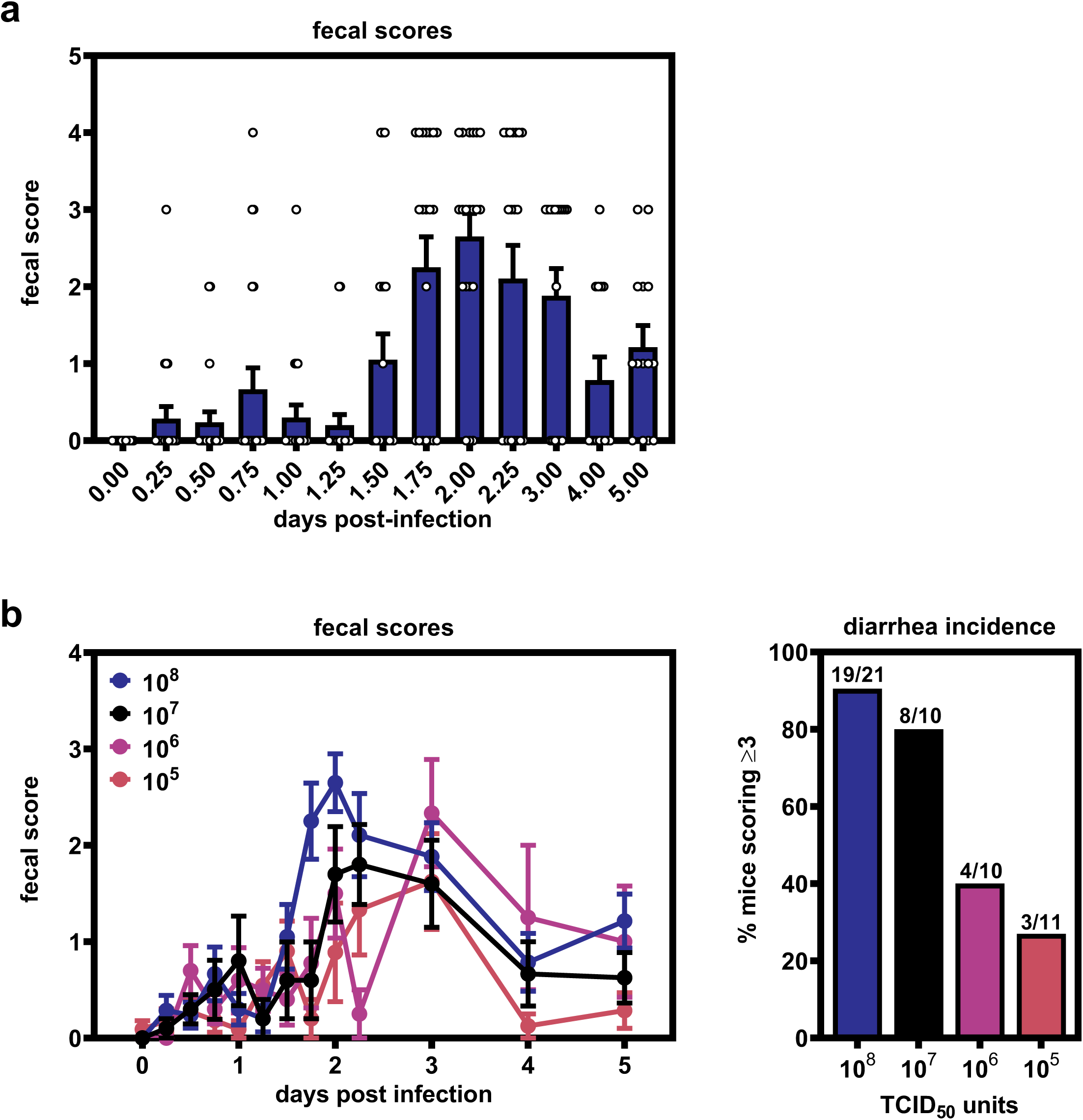
MNV-1-induced diarrhea in neonatal mice is dose-dependent. **a)** The data for MNV-1-infected BALB/c pups presented in Fig. 1c is shown here as a bar graph with scores for individual mice represented by dots. **b)** Groups of 3-day old BALB/c pups were infected with 10^8^ (*n* = 21), 10^7^ (*n =* 10), 10^6^ (*n =* 10), or 10^5^ (*n* = 11) TCID_50_ units of MNV-1 by oral gavage and monitored for fecal consistency by palpating their abdomens. Error bars denote standard errors of mean in all figures.

**Supplementary Figure 2.**
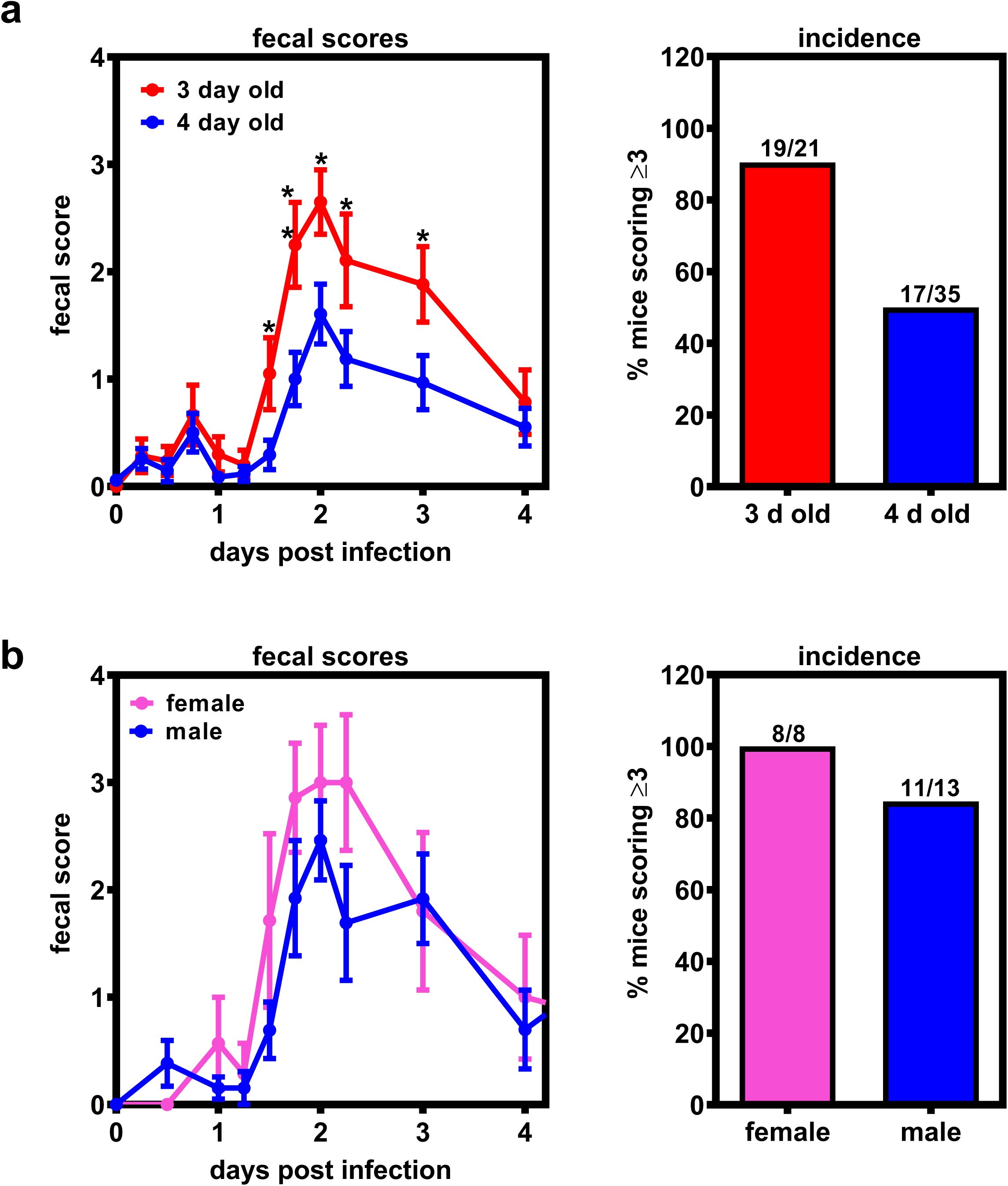
Host age and sex influence MNV disease severity in neonatal mice. **a)** Data stratified by age from groups of 3- and 4-day old BALB/c pups (*n =* 21 for 3-day old; *n* = 35 for 4-day old) infected with 10^8^ TCID_50_ units of MNV-1 by oral gavage and monitored for fecal inconsistency. **b)** Data stratified by sex (*n =* 8 for female; *n =* 13 for male) from groups of 3-day old BALB/c pups infected with 10^8^ TCID_50_ units of MNV-1 by oral gavage and monitored for fecal inconsistency. Error bars denote standard errors of mean in all figures. *P* values were determined using two-way ANOVA with corrections for multiple comparisons. (*P < 0.05, **P < 0.01, ***P < 0.001).

**Supplementary Figure 3.**
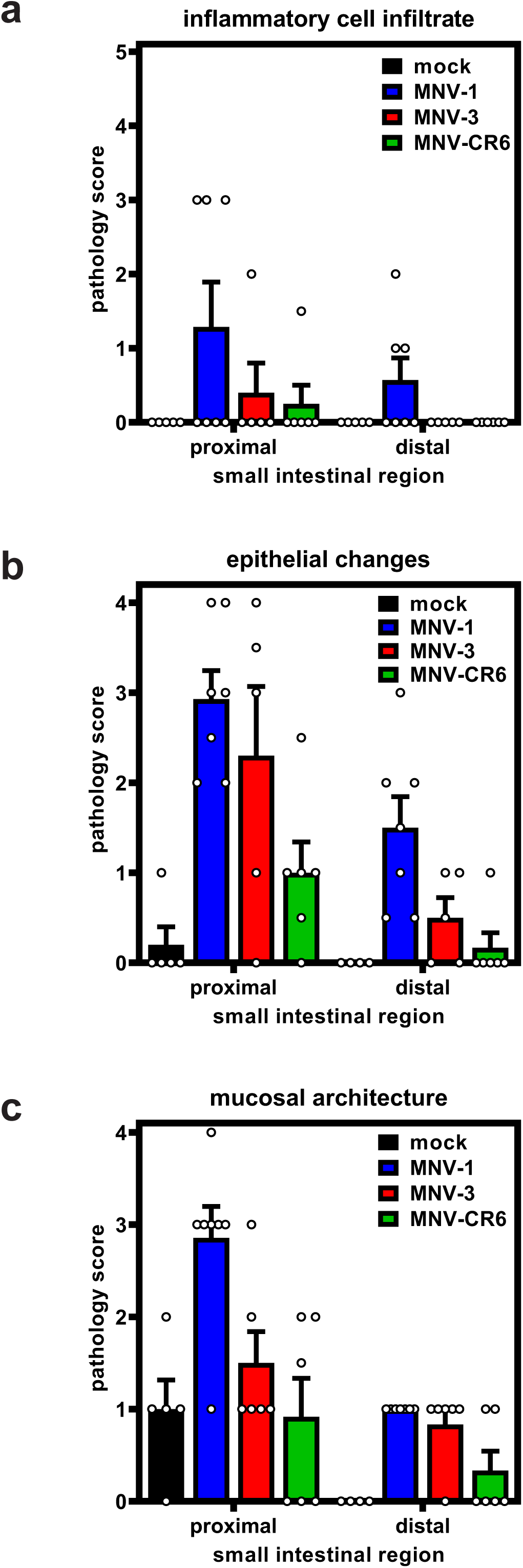
Pathology scores for each scoring criteria. Small intestinal sections collected from neonates infected with 10^8^ TCID_50_ units of MNV-1, MNV-3, MNV-CR6, or mock inoculum at 2 dpi (*n =* 5 for mock, *n =* 7 for MNV-1, *n =* 5 for MNV-3, *n =* 6 for MNV-CR6) were stained with hematoxylin and eosin. Sections were scored blindly by an animal veterinarian for pathological changes according to three criteria, as defined in the methods.

## REFERENCEES

1. Koo, H.L. et al. J. Pediatr. Infect. Dis. Soc. 2, 57–60 (2013).

2. Patel, M.M. Emerg. Infect. Dis. 14, 1224–1231 (2008).

3. Ahmed, S.M. et al. Lancet Infect. Dis. 14, 725–730 (2014).

4. Cheetham, S. et al. J Virol 80, 10372–10381 (2006).

5. Seo, D.J. et al. J. Med. Virol. 90, 655–662 (2018).

6. Souza, M., Azevedo, M.S.P., Jung, K., Cheetham, S. & Saif, L.J. J Virol 82, 1777–1786 (2008).

7. Karst, S.M., Wobus, C.E., Lay, M., Davidson, J. & Virgin, H.W. Science 299, 1575–1578 (2003).

8. Grau, K.R. et al. Nat. Microbiol. 2, 1586 (2017).

9. Hsu, C.C., Riley, L.K., Wills, H.M. & Livingston, R.S. Comp. Med. 56, 247–251 (2006).

10. Arias, A., Bailey, D., Chaudhry, Y. & Goodfellow, I.G. J. Gen. Virol. 93, 1432–1441 (2012).

11. Thackray, L.B. et al. J Virol 81, 10460–10473 (2007).

12. Wilen, C.B. et al. Science 360, 204–208 (2018).

13. Mumphrey, S.M. et al. J Virol 81, 3251–3263 (2007).

14. Kahan, S.M. et al. Virology 421, 202–210 (2011).

15. Zhu, S. et al. J. Virol. 90, 2858–2867 (2016).

16. Rocha-Pereira, J., Kolawole, A.O., Verbeken, E., Wobus, C.E. & Neyts, J. Antiviral Res. 132, 76–84 (2016).

17. Strong, D.W., Thackray, L.B., Smith, T.J. & Virgin, H.W. J. Virol. 86, 2950–2958 (2012).

18. Feng, N., Franco, M.A. & Greenberg, H.B. Adv. Exp. Med. Biol. 412, 233–240 (1997).

19. Ramig, R.F. Microb. Pathog. 4, 189–202 (1988).

20. Du, J. et al. Virus Res. 228, 134–140 (2017).

21. Cannon, J.L., Lopman, B.A., Payne, D.C. & Vinjé, J. Clin. Infect. Dis. 69, 357–365 (2019).

22. Menon, V.K. et al. PLOS ONE 11, e0157007 (2016).

23. Rouhani, S. et al. Clin. Infect. Dis. Off. Publ. Infect. Dis. Soc. Am. 62, 1210–1217 (2016).

24. Agus, S.G., Dolin, R., Wyatt, R.G., Tousimis, A.J. & Northrup, R.S. Ann. Intern. Med. 79, 18–25 (1973).

25. Schreiber, D.S., Blacklow, N.R. & Trier, J.S. J. Infect. Dis. 129, 705–708 (1974).

26. Parrino, T.A., Schreiber, D.S., Trier, J.S., Kapikian, A.Z. & Blacklow, N.R. N. Engl. J. Med. 297, 86–89 (1977).

27. Schreiber, D.S., Blacklow, N.R. & Trier, J.S. N. Engl. J. Med. 288, 1318–1323 (1973).

28. Dolin, R., Levy, A.G., Wyatt, R.G., Thornhill, T.S. & Gardner, J.D. Am. J. Med. 59, 761–768 (1975).

29. Troeger, H. et al. Gut 58, 1070–1077 (2009).

30. Seamons, A. et al. Am. J. Pathol. 188, 1536–1554 (2018).

31. González-Hernández, M.B., Perry, J.W. & Wobus, C.E. Bio-Protoc. 3, (2013).

32. Perry, J.W. & Wobus, C.E. J. Virol. 84, 6163–6176 (2010).

33. Ball, J.M., Tian, P., Zeng, C.Q., Morris, A.P. & Estes, M.K. Science 272, 101–104 (1996).

34. Lundgren, O. et al. Science 287, 491–495 (2000).

35. Istrate, C., Hagbom, M., Vikström, E., Magnusson, K.-E. & Svensson, L. J. Virol. 88, 3161–3169 (2014).

36. Zhu, S. et al. PLoS Pathog 9, e1003592 (2013).

37. Gonzalez-Hernandez, M.B. et al. J. Virol. 88, 6934–6943 (2014).

38. Zhu, S. et al. PLoS Pathog. 9, e1003592 (2013).

39. Erben, U. et al. Int. J. Clin. Exp. Pathol. 7, 4557–4576 (2014).

